# Membrane-mediated interactions between hinge-like particles

**DOI:** 10.1101/2022.01.19.476938

**Authors:** Bing Li, Steven M. Abel

## Abstract

Adsorption of nanoparticles on a membrane can give rise to interactions between particles, mediated by membrane deformations, that play an important role in self-assembly and membrane remodeling. Previous theoretical and experimental research has focused on nanoparticles with fixed shapes, such as spherical, rod-like, and curved nanoparticles. Recently, hinge-like DNA origami nanostructures have been designed with tunable mechanical properties. Inspired by this, we investigate the equilibrium properties of hinge-like particles adsorbed on an elastic membrane using Monte Carlo and umbrella sampling simulations. The configurations of an isolated particle are influenced by competition between bending energies of the membrane and the particle, which can be controlled by changing adsorption strength and hinge stiffness. When two adsorbed particles interact, they effectively repel one another when the strength of adhesion to the membrane is weak. However, a strong adhesive interaction induces an effective attraction between the particles, which drives their aggregation. The configurations of the aggregate can be tuned by adjusting the hinge stiffness: Tip-to-tip aggregation occurs for flexible hinges, whereas tip-to-middle aggregation also occurs for stiffer hinges. Our results highlight the potential for using the mechanical features of deformable nanoparticles to influence their self-assembly when the particles and membrane mutually influence one another.

## 1 Introduction

Membranes can be transformed from one state to another by the adsorption of nanoparticles. Membrane-adsorbed nanoparticles commonly induce local curvature of the membrane, and the local curvature induced by one nanoparticle can influence other nanoparticles in the vicinity. As a result, the nanoparticles can experience attractive interactions that favor their self-assembly and promote large-scale membrane deformations.^1–5^

Membrane-mediated interactions between nanoparticles arise largely as a result of the overall bending energy of the membrane. It has been revealed both experimentally^6–8^ and theoretically^2,3,9–11^ that the bending energy depends on the distances between nanoparticles. For example, when curvature-inducing colloidal spheres are adsorbed on planar lipid bilayers, the spherical nanoparticles experience an attractive force when they come close to each other.^9^ The net force between the nanoparticles results from the change of the bending energy as a function of the distance between them. For spherical nanoparticles adsorbed on planar fluid membranes, linear aggregates of nanoparticles have been observed in simulations and explained by attractive three-particle interactions.^2^ For spherical nanoparticles on the inside^3^ or outside^10^ of spherical vesicles, it has been found that the nanoparticles also experience a mutual attraction and form linear aggregates enclosed by membrane tubules, which protrude out of or into the vesicles.

The bending energy of membranes also depends on the relative orientations of adsorbed, anisotropic nanoparticles.^11–13^ For instance, rod-like fd viruses adsorbed onto a cationic lipid bilayer tend to form tip-to-tip linear aggregates at lower densities.^13^ This lowers the bending energy around their tips by connecting the ends and reducing the highly curved area of the underlying membrane. Computer simulations further showed that tip-to-tip aggregation is favored for soft membranes, while side-by-side contact is preferred for stiffer membranes.^1^ Additionally, the shape of a nanoparticle can affect the cellular uptake of particles.^14,15^

The assembly of curved nanoparticles on membranes has been of particular interest because of their similarity to BAR (Bin/Amphiphysin/Rvs) proteins.^16–22^ BAR proteins can be modelled as curved rods, and they have been shown to both sense and generate local curvature of membranes. Coarse-grained molecular dynamics simulations of BAR proteins adsorbed onto membranes showed that tip-to-tip aggregation is favorable at strong adhesion strength.^4,23^ For tensionless membranes, tip-to-tip aggregation of proteins is predominant, but large membrane tension favors side-by-side aggregation to maximize the contact surface of proteins. Other simulations of rigid, curved nanoparticles showed that nanoparticles adsorbed on membrane vesicles assemble into two types of aggregates: side-to-side and tip-to-tip, depending on the adhesion strength.^12^

DNA origami nanostructures have significant promise for modifying properties of membranes and remodeling their shape.^24–28^ Curved DNA origami objects, designed to mimic structural and functional features of BAR domain proteins, can induce curvature of lipid bilayers and reproduce features of membrane-sculpting proteins.^29^ DNA origami curls on membranes can polymerize into nanosprings through linker strands at the ends of the curls, inducing membrane tubulation.^30^. Recently, considerable progress has been made in designing deformable DNA origami nanostructures with controllable mechanical features. Work by the Castro group first introduced deformable, hinge-like DNA origami nanostructures in which the preferred angle of the hinge and its mechanical properties can be tuned by relatively small adjustments in the design.^31,32^ Such structures can be used as a basis for the design of mechanically functional DNA origami devices and materials, and it yet remains to explore their behavior when interacting with a membrane.

In previous work, we explored the adsorption of semiflexible polymers on membrane vesicles and revealed a complex interplay between deformations of the membrane and polymer.^33^ This interplay has motivated our interest in the study of deformable nanostructures adsorbed to membranes because such systems have the potential to exhibit rich interactions and collective behavior. In this study, we use computer simulations to study equilibrium properties of hinge-like particles adsorbed on a deformable membrane. We first describe the model and simulation details. We then study how the configurations of a single adsorbed particle are impacted by the hinge stiffness and the adsorption strength. We investigate interactions between two adsorbed particles and characterize their dependence on the adsorption strength and hinge stiffness. Umbrella sampling is then used to determine the potential of mean force as a function of the distance r between the centers of mass of two particles, first for flexible hinges and then for stiffer hinges. We conclude by discussing our results in the context of the self-assembly of deformable nanoparticles.

## 2 Model and simulation details

The membrane is represented as an infinitely thin elastic surface consisting of *M* = 418 spherical hard beads of diameter *l*_*mem*_ = *σ* connected by bonds to form a triangulated and self-avoiding network. We consider a planar membrane with periodic boundary conditions. The bond length can range from *σ* to 1.67*σ*. The membrane bending energy is given by^34–36^

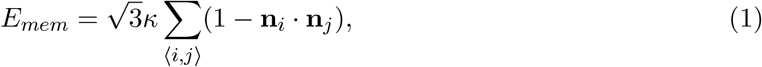

where *κ* is the bending rigidity of the membrane, the sum is over all triangles *i* and *j* sharing an edge, and **n**_*i*_ denotes the normal vector to triangle *i*. In our work, *κ* is set 10 *k*_B_*T*. We consider a fluid membrane in which the connectivity of the membrane is dynamically rearranged to simulate the fluidity of the membrane.^33,36,37^ The surface tension *γ* dictates the cost associated with area changes, *E*_*γ*_ = *γA*, where *A* is the total surface area. We set *γ* = 1 *k*_B_*T/σ*^2^ in our simulations, which corresponds to the surface tension of the order of 10^−2^ to 10^−3^ pN*/*nm.^2^ The minimum membrane area is *A* ≈ 643*σ*^2^.

The hinge-like particle is represented by two connected rods (see Fig. 1, inset). Each rod consists of three hard sphere beads of diameter *l*_*r*_ = 2*σ* connected by fixed-length bonds of length 2.02*σ*. The rods share a common bead at their ends, and the angle between the two arms of the hinge is denoted by *θ*. The bending energy of the hinge is^33^

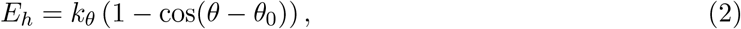

where *k*_*θ*_ is the hinge stiffness and *θ*_0_ is the preferred hinge angle. For this work, we let *θ*_0_ = *π/*2. The surface fraction covered by a single hinge on the membrane is *ρ* = 0.03.

**Figure 1:**
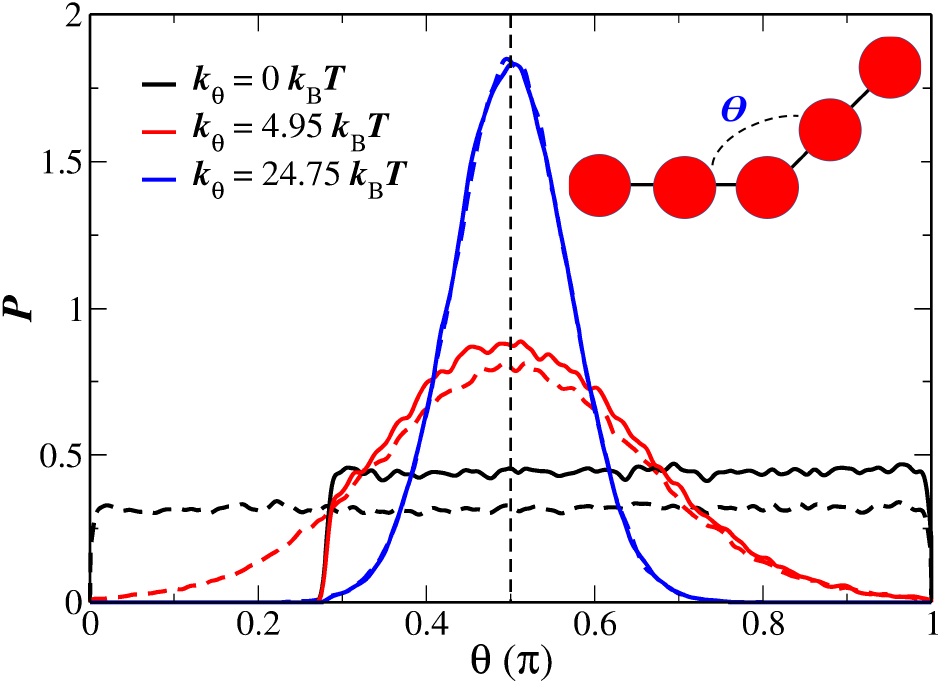
The probability density (*P*) of the hinge angle (*θ*) for different values of the hinge stiffness without a membrane. The dashed lines correspond to ideal particles without hard-sphere interactions between beads of the hinge. The solid lines correspond to particles with hard-sphere interactions. Inset is a schematic of the hinge-like particle.

The particle-to-membrane adhesion is modeled via a generic power-law potential between particle and membrane beads. The interaction between bead *i* of the particle and bead *j* of the membrane is given by^3^

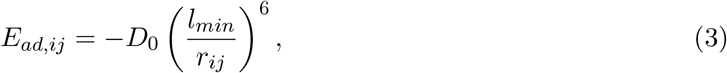

where *l*_*min*_ = (*l*_*mem*_ + *l*_*r*_)*/*2, *r*_*ij*_ is the distance between the beads, and a cutoff is imposed at *r*_*cut*_ = 1.5 *l*_*min*_. The total adhesion energy *E*_*ad*_ is the sum over all pairs of *i* and *j*. We consider three strengths of the adhesion potential: *D*_0_ = 5, 10, and 15 *k*_B_*T*. The total energy of the system is *E*_total_ = *E*_*mem*_ + *E*_*γ*_ + *E*_*h*_ + *E*_*ad*_.

We use Metropolis Monte Carlo computer simulations to sample configurations of the system at thermal equilibrium. For membranes, there are two types of trial Monte Carlo (MC) moves: single-particle displacement moves and bond-flip moves. The details are presented in previous work.^33,38^ For nanoparticles, we use a pivot move,^39,40^ where one rod of nanoparticles is randomly selected and rotated by a random angle around the axis through the hinge bead with random orientation. All trial moves are accepted or rejected according to the standard Metropolis criterion, and the simulations satisfy detailed balance. Each Monte Carlo step (MCS) consists of *M* attempted displacement moves, *M* attempted bond-flip moves, and 100 attempted pivot moves.

We first equilibrate the system for 1 × 10^6^ MCS with nanoparticles and the membrane well separated so that they equilibrate independently. We then place the nanoparticles near one side of the membrane surface such that the minimum distance between beads of nanoparticles and the membrane is *r*_*cut*_. For adsorption strengths of *D*_0_ = 10 and 15 *k*_B_*T*, we employ a simulated annealing method to obtain reliable sampling.^33^ The value of *D*_0_ is increased from *D*_0_ = 5 *k*_B_*T* to the target value with an increment *δϵ* = 0.2 *k*_B_*T*. At each increment, we relax the system for 1 × 10^6^ MCS. Upon reaching the target value, we relax the system for an additional 5 × 10^6^ MCS before collecting the data. We perform 1 × 10^7^ MCS in each collection run and store the configuration every 5 × 10^3^ MCS. Ten independent simulation trajectories are generated for each set of conditions.

We also employ umbrella sampling simulations to calculate the potential of mean force (PMF) as a function of the distance between two particles adsorbed on the membrane.^40–44^ We apply a harmonic bias potential, 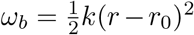, between the two particles. Here, the reaction coordinate *r* is the distance between the centers of mass of the two particles. The harmonic spring constant *k* is set to the value of 200*k*_B_*T/σ*^2^ to allow sampling of features across a wide range of *r*. To generate overlapping windows, we divide the reaction coordinate *r* ∈ [2*σ*, 12.68*σ*] into 39 windows with different *r*_0_. The maximum distance, *r*_max_ = 12.68*σ*, is set to the half of the simulation box size. We use the weighted histogram analysis method (WHAM) to obtain the unbiased probability distribution, from which we calculate the free energy difference of the system relative to the free energy at *r*_*max*_: Δ*F* = *F* (*r*) − *F* (*r*_max_).^44^ We also calculate the membrane bending energy, adsorption energy, and total energy by averaging configurations falling into the neighbourhood of each window center, *r* ∈ [*r*_0_ − *d, r*_0_ + *d*], where *δ* = 0.1*σ* is used to generate sufficient sampling.

## 3 Results and discussion

For fixed-shape nanoparticles such as spherical nanoparticles, the shape of a membrane around a single particle can be tuned by adjusting the strength of the adhesive interaction. The nanoparticles can be slightly, partially, or totally wrapped by the membrane depending on the strength of attraction, which must be sufficiently strong to compensate for the cost of bending the membrane.^45–48^ Unlike rigid particles, hinge-like nanoparticles do not have a fixed shape, and it remains to explore the equilibrium properties of adsorbed hinges. The hinge stiffness introduces a new energy into the system that can lead to competition between membrane bending and hinge deformations.

### 3.1 Single particle: Interplay of adhesion strength, membrane deformations, and hinge stiffness

We first study the equilibrium properties of an isolated, hinge-like nanoparticle. Figure 1 shows the distribution of hinge angles (*θ*) sampled without a membrane present. Results are shown for an idealized hinge in which the the hard-sphere interactions between the beads of the hinge are neglected (dashed lines) and for the hinge with hard-sphere interactions, as described in the methods (solid lines). For a fully flexible hinge (*k*_*θ*_ = 0), the distribution of angles is uniform. With hard-sphere interactions, there is a cutoff at *θ* ≈ 0.28*π* because smaller angles would lead to overlap of beads on the different arms of the hinge. For nonzero hinge stiffness, the angle distribution has a peak at *θ* = *π/*2, with a variance that decreases with increasing *k*_*θ*_. The cutoff due to hard-sphere interactions impacts the hinge with *k*_*θ*_ = 4.95 *k*_B_*T* but has minimal impact on the stiffest hinge. As expected, the distribution of angles sampled by the particle can be tuned by changing the stiffness. Figure 2 shows the probability density of the hinge angle for a single hinge adhered to the surface of a membrane. We consider three values of the adsorption strength (*D*_0_ = 5, 10, and 15 *k*_B_*T*) and three values of the hinge stiffness (*k*_*θ*_ = 0, 4.95 and 24.75 *k*_B_*T*). At the weakest adsorption strength (*D*_0_ = 5 *k*_B_*T*, Fig. 2a), the distribution of the hinge angle *θ* is similar to the distribution for particles without a membrane. Increasing the adsorption strength to *D*_0_ = 10 *k*_B_*T* (Fig. 2b), the distribution of angles with *k*_*θ*_ = 0 changes substantially, with a peak at *θ* ≈ 0.9*π*. There is a modest change in the peak location and shape of the distribution for *k*_*θ*_ = 4.95 *k*_B_*T* and little change in the distribution for *k*_*θ*_ = 24.75 *k*_B_*T*. At the strongest adsorption strength (*D*_0_ = 15 *k*_B_*T*, Fig. 2c), all of the hinges exhibit some degree of straightening: The distributions shift toward larger angles, although the peak occurs at smaller angles for stiffer hinges.

**Figure 2:**
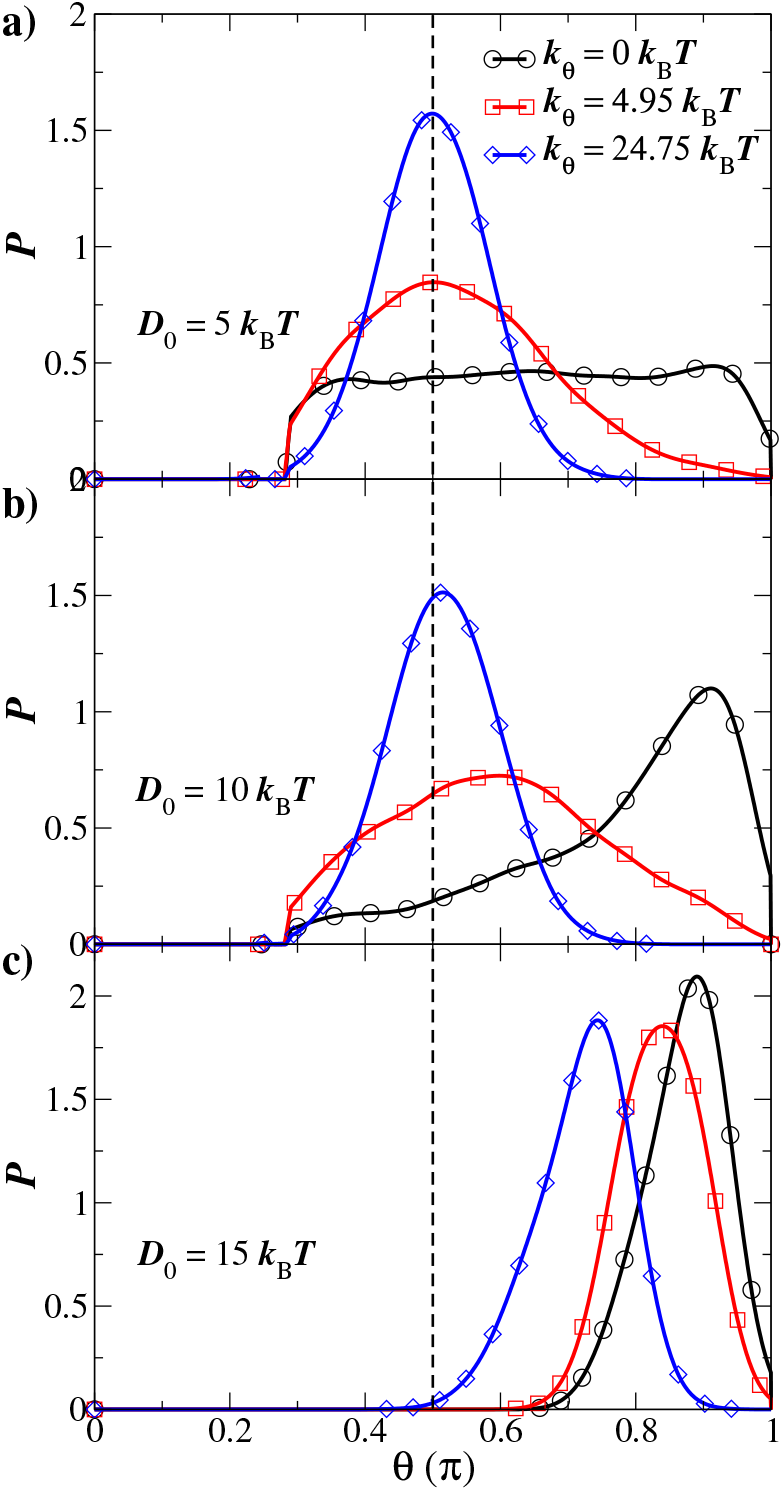
The probability density (*P*) of the hinge angle (*θ*) at different values of the adsorption strength (*D*_0_) and hinge stiffness (*k*_*θ*_). The preferred hinge angle (*θ* = *π/*2) is denoted by the vertical dashed line.

The results in Fig. 2 can be understood in terms of the wrapping of the membrane around the particles. At the weakest adsorption strength (*D*_0_ = 5 *k*_B_*T*), the membrane is minimally deformed and thus has little influence on the configurations of particles. At *D*_0_ = 10 *k*_B_*T*, the particles are slightly wrapped by membrane. The shape of flexible particles (*k*_*θ*_ = 0) tends to straighten so as to decrease the bending energy of the membrane. However, for the stiffer hinges, the configurations are dominated by the bending energy of the particles. At *D*_0_ = 15 *k*_B_*T*, the particles are partially wrapped by the membrane and the membrane bending energy becomes relevant for all hinge stiffnesses. This results in the shift toward larger angles for *k*_*θ*_ = 4.95 and 24.75 *k*_B_*T*. In this regime, interplay between the bending energies of the particle and the membrane becomes relevant. Note that the position of the peak for *k*_*θ*_ = 0 is not at *θ* = *π*, where the hinge is totally straight. We suggest that it is because a slight bend of the hinge enhances the adhesion energy because some membrane beads can interact with both rods of the nanoparticle.

At an even stronger adsorption strength (*D*_0_ = 20 *k*_B_*T*), we find the hinge is totally wrapped by the membrane in a bud-like state. The two arms of the hinge are squeezed toward each other and the peak of the angle distribution occurs at *θ* ≈ 0.33*π*.

### 3.2 Two particles: Effective interactions of flexible hinges

The results in Fig. 2 demonstrate membrane-mediated changes to configurations of a single hinge. Adhesion-induced membrane curvature can also lead to interactions between two particles. Here, we study the equilibrium properties of two hinge-like particles adsorbed to a membrane.

We first consider the case of flexible hinges (*k*_*θ*_ = 0), for which the behavior of the system is governed primarily by the interplay of the membrane bending energy and the adsorption energy. With *D*_0_ = 5 *k*_B_*T*, the membrane has minimal influence on the particles. We thus consider the larger values of the adsorption strength, *D*_0_ = 10 and 15 *k*_B_*T*. In our unbiased simulations, at *D*_0_ = 10 *k*_B_*T*, the two particles remain separated, suggesting an effective repulsion at short distances. In contrast, at *D*_0_ = 15 *k*_B_*T*, the two particles form a tip-to-tip aggregate by coming into contact at their ends. Representative snapshots are shown in the upper panel of Fig. 3. The simulations indicate that the two particles experience an effective attraction at *D*_0_ = 15 *k*_B_*T*.

**Figure 3:**
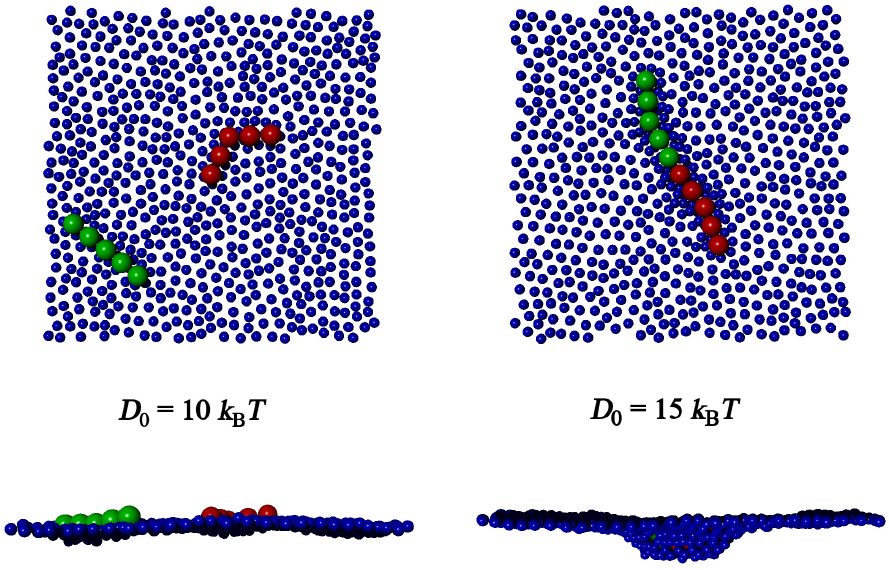
Typical equilibrium configurations of two flexible hinges (*k*_*θ*_ = 0) adsorbed to a membrane viewed from the top (upper panel) and side (lower panel). Two strengths of the attractive potential are shown: *D*_0_ = 10 *k*_B_*T* (left) and 15 *k*_B_*T* (right).

As expected from earlier theoretical and computational work, particles can be partially wrapped by a membrane when the adhesive interaction is sufficiently strong. The lower panel of Fig. 3 shows the degree of wrapping with *D*_0_ = 10 and 15 *k*_B_*T*. The stronger attraction induces a larger membrane deformation and leads to extended particles that are almost straight.

To quantify the effective interactions between hinges, we characterize the potential of mean force (PMF) as a function of the distance (*r*) between the centers of mass of two particles (Fig. 4a). For these calculations, we use a larger system (*M* = 672) to reduce the effects of system size and to characterize larger separation distances. At *D*_0_ = 10 *k*_B_*T*, the PMF shows a purely repulsive interaction between two particles. In this regime, the total energy is approximately constant (Fig. 4b), and the effective repulsion is entropic in nature. When the hinges approach one another, the entropy decreases because there is a decrease in configurational entropy.^44^

**Figure 4:**
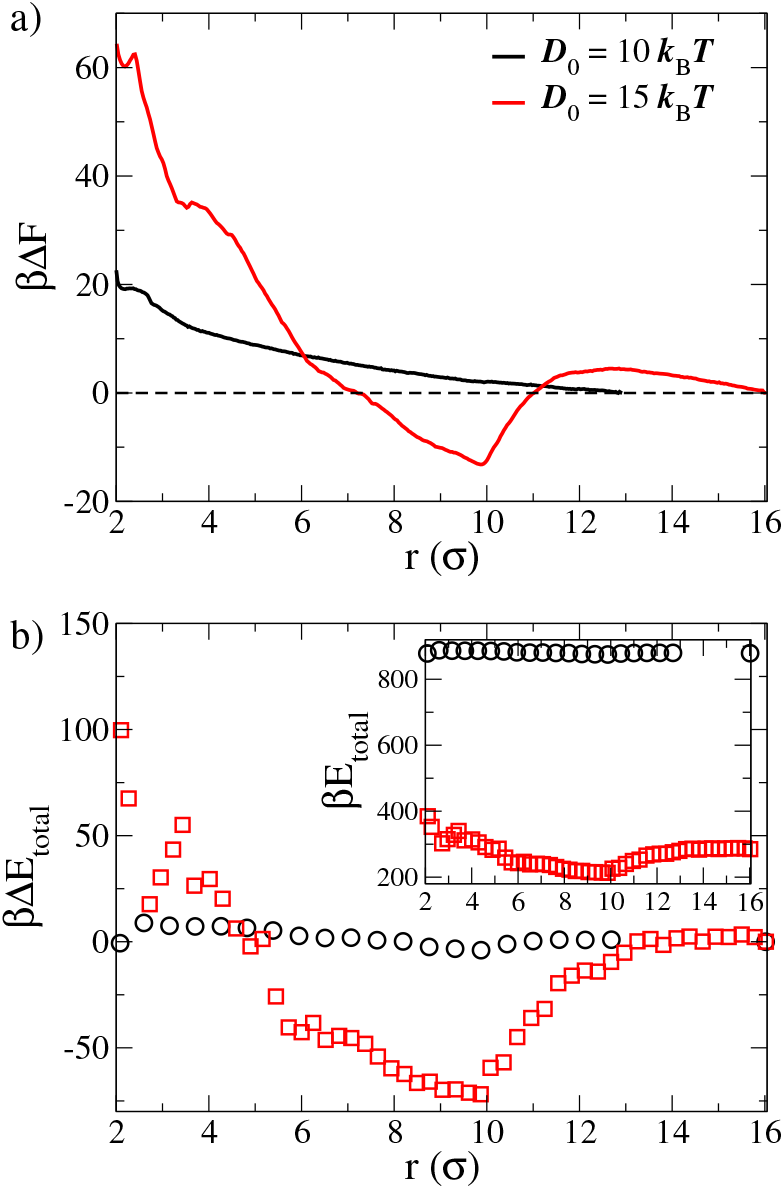
(a) Potential of mean force (PMF), Δ*F*, as a function of the distance (*r*) between the centers of mass of two hinges. (b) Total energy, *E*_total_, of the system as a function of *r*. Two adsorption strengths, *D*_0_ = 10 and 15 *k*_B_*T*, are studied with *k*_*θ*_ = 0. Δ*E* denotes the change in energy relative to the energy at *r*_max_. *β* = 1*/k*_B_*T*.

At *D*_0_ = 15 *k*_B_*T*, the PMF exhibits an attractive well. At the largest distances considered, the effective interaction is slightly repulsive, but the particles begin to experience an attractive interaction when the distance decreases to *r* ≈ 12.5*σ*. The minimum occurs at *r* ≈ 10*σ*, which is associated with tip-to-tip aggregation of the particles (see Fig. 3). The energy barrier at large distances is approximately 4.5 *k*_B_*T* and the minimum of the PMF is approximately −14 *k*_B_*T*. At smaller distances, the interaction is repulsive. The shape near the minimum is similar to that of the total energy (Fig. 4b), indicating that the PMF is significantly influenced by the total energy. For *r <* 6*σ*, the value of free energy difference with *D*_0_ = 15 *k*_B_*T* is larger than one with *D*_0_ = 10 *k*_B_*T*. Thus, even though the stronger adsorption results in an effective attraction favoring end-to-end alignment, it is more costly to bring the centers of mass into close contact (e.g., side-by-side) compared with the weaker adsorption.

Figure 5 further explores the adsorption energy and membrane bending energy as a function of the distance between hinges. With *D*_0_ = 10 *k*_B_*T*, both the adsorption energy and bending energy are relatively constant. This is consistent with the adsorbed particles inducing only slight deformations of the membrane. The adsorption energy increases modestly for *r* ≲ 6*σ*, but the effect is offset by a decrease in the membrane bending energy. In this regime, the membrane incurs a smaller bending penalty to accommodate the two particles in close proximity, but it has less contact area with the particles (fewer membrane contacts).

**Figure 5:**
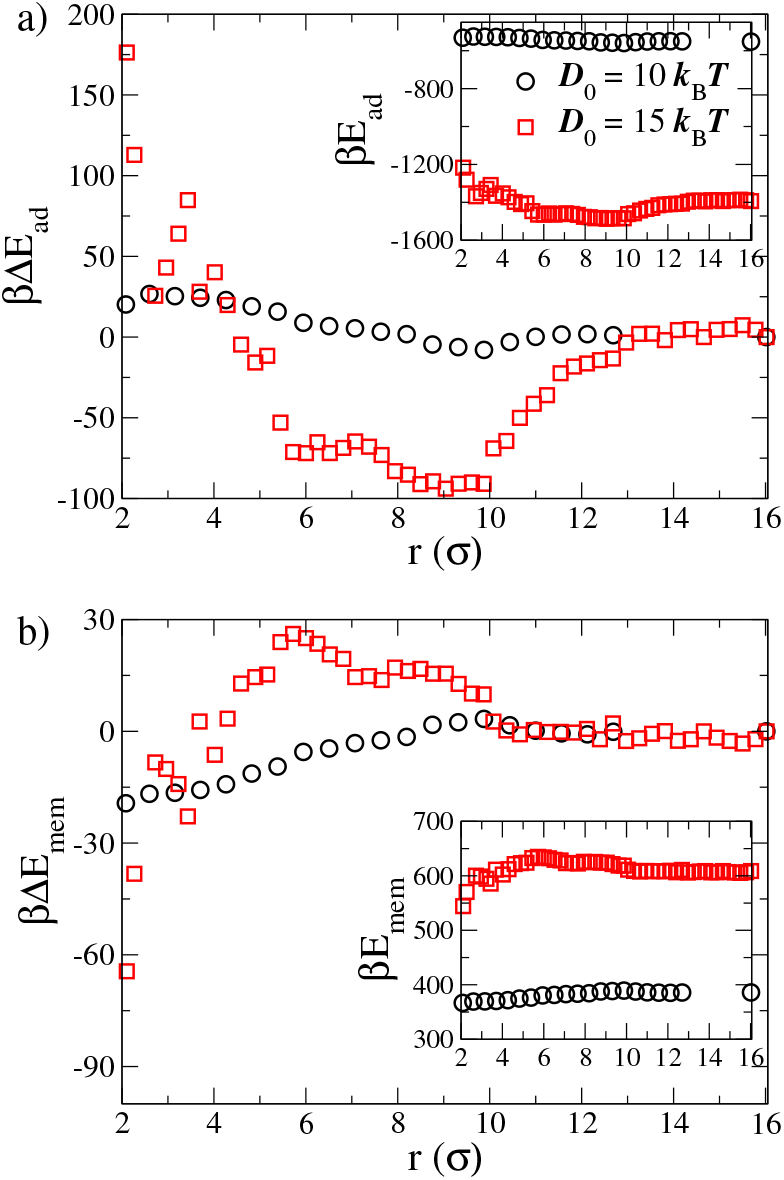
(a) Adsorption energy, *E*_*ad*_, as a function of the distance (*r*) between the centers of mass of two nanoparticles. (b) Membrane bending energy, *E*_*mem*_, as a function of *r*. Two adsorption strengths, *D*_0_ = 10 and 15 *k*_B_*T*, are studied with *k*_*θ*_ = 0. Δ*E* denotes the change in energy relative to the energy at *r*_max_. *β* = 1*/k*_B_*T*.

With *D*_0_ = 15 *k*_B_*T*, there are much more substantial changes in both the adsorption energy and membrane bending energy. Comparing with the total energy (Fig. 4b) reveals that the adsorption energy has a similar shape and makes a larger contribution than membrane bending energy to the change in total energy. In the region of *r* near the attractive well of the PMF, Δ*E*_*ad*_ *<* 0 indicates an increase in contact between the membrane and particles. This is coupled with an increase in the bending energy (Δ*E*_*mem*_ *>* 0). This suggests that the particles adopt an end-to-end configuration because a shared deformation enables more complete wrapping of the particles at a modest increase to the bending energy. In isolation, increased wrapping would be more energetically costly.

Smaller values of *r* behave similarly to *D*_0_ = 10 *k*_B_*T* but with larger energy scales. Figure 6 shows sample configurations when *r <* 4*σ*. With *r* = 3.7*σ*, the particles adopt configurations that would require large bending energies of the membrane to wrap all sides of the nanoparticles. Other configurations are in side-to-side contact (e.g., those at *r* = 2.65*σ* and 2*σ*). This leads to a decrease of the number of bound membrane beads because of excluded volume interactions that prevent the membrane from wrapping the inner surface of the particles. Physically, when the two particles are close together, the membrane adopts a less pronounced deformation when wrapping the two particles together. However, this occurs at a cost to the amount of surface area in contact.

**Figure 6:**
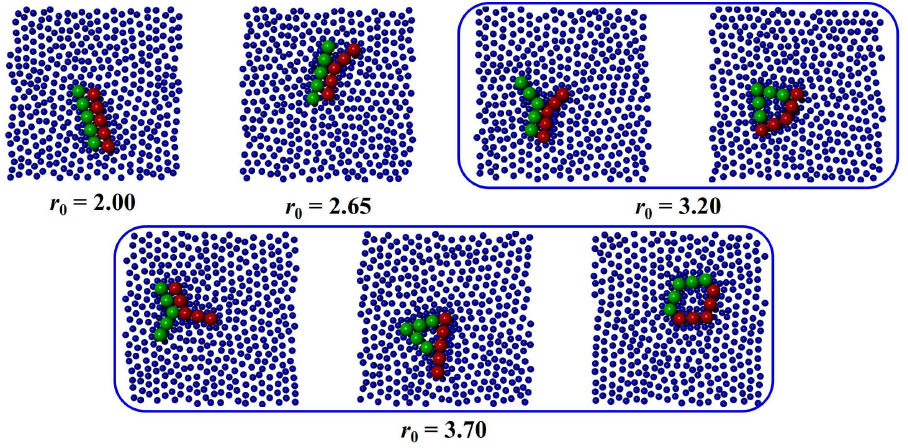
Snapshots of configurations of two particles at various separation distances (*r*_0_) with *D*_0_ = 15 *k*_B_*T* and *k*_*θ*_ = 0.

### 3.3 Two particles: Impact of hinge stiffness

We next consider the influence of the hinge stiffness on the effective interaction between two adsorbed particles. For a single adsorbed particle, the equilibrium configurations are determined by the interplay of the bending energies of both the membrane and the particle. Additionally, we showed that two strongly adsorbed flexible hinges experience an effective attraction that favors tip-to-tip aggregation with straightened hinge configurations. However, with a nonzero hinge stiffness, this configuration will be less favorable, making it interesting to study how the hinge stiffness impacts the effective interaction between two particles.

As before, with *D*_0_ = 10 *k*_B_*T*, the membrane deformation is modest and the effective interaction between the two particles is dominated by entropic considerations. This leads to an effective repulsion of the two particles for *k*_*θ*_ = 4.95 and 24.75 *k*_B_*T*. In the following, we focus on the stronger adsorption strength (*D*_0_ = 15 *k*_B_*T*), which leads to larger deformations and an effective attraction when *k*_*θ*_ = 0.

Figure 7 shows the PMF as a function of *r* with *D*_0_ = 15 *k*_B_*T* and *k*_*θ*_ = 24.75 *k*_B_*T*. As for *k*_*θ*_ = 0, there is an attractive minimum with a small energy barrier at large distances. However, both the depth and location of the minimum are different. When the hinge stiffness increases from *k*_*θ*_ = 0 to 24.75 *k*_B_*T*, the minimum value of the free energy increases, indicating that the strength of the attraction decreases with an increase of the hinge stiffness. The location of the minimum of the free energy decreases from *r* ≈ 10*σ* to *r* ≈ 9*σ*.

**Figure 7:**
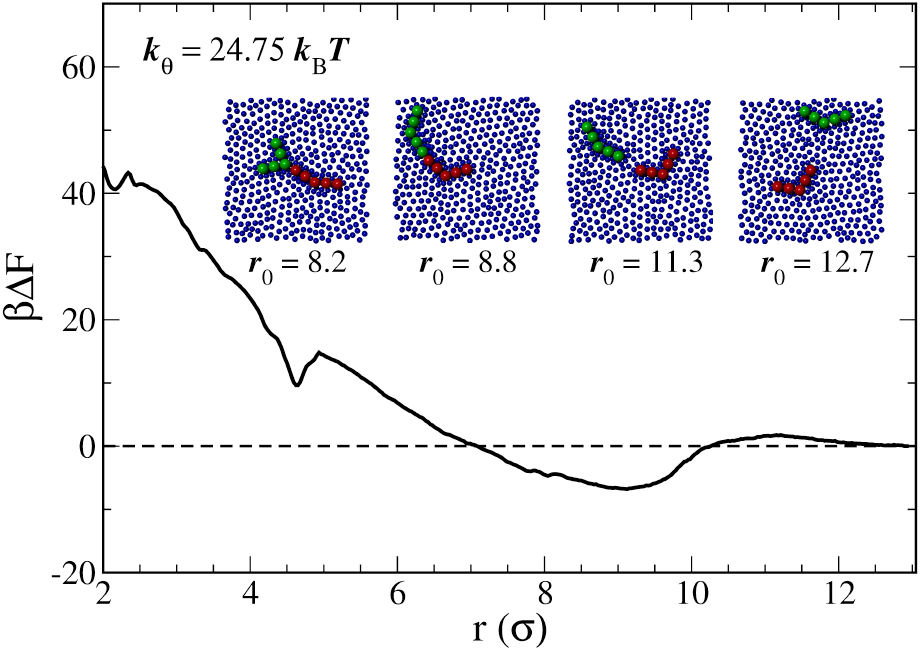
Potential of mean force, Δ*F*, as a function of the distance (*r*) between the centers of mass of two hinges with *D*_0_ = 15 *k*_B_*T* and *k*_*θ*_ = 24.75 *k*_B_*T*. *β* = 1*/k*_B_*T*. Inset: Snapshots of typical configurations at four distances: *r* = 12.7*σ* (far apart and non-interacting), *r* = 11.3*σ* (near the peak of the small energy barrier), and *r* = 8.8*σ* and 8.2*σ* (near the minimum of the attractive energy well).

The changes in the PMF are a result of the hinge stiffness resulting in less favorable configurations compared to the flexible hinge. The minimum is at smaller values of *r* because the hinges prefer to remain bent, thus bringing their centers of mass closer together. For *k*_*θ*_ = 24.75 *k*_B_*T*, in the basin of attraction near *r* ≈ 9*σ* two classes of configurations are commonly observed: tip-to-tip aggregation of bent conformations and tip-to-middle aggregation (Fig. 7, inset). Thus, increasing the hinge stiffness results in a previously unobserved equilibrium configuration (tip-to-middle).

It is also interesting to note that the PMF has an apparent local minimum at *r* = 4.6*σ*, which is in the strongly repulsive regime. At this distance, the nanoparticles form a square-like aggregate by connecting both ends of each particle (similar to the right snapshot of the lower panel of Fig. 6).

## 4 Conclusions

Adsorption of particles on a membrane can give rise to effective repulsive and attractive interactions between particles that are mediated by deformations of the membrane. Most studies to date have focused on particles of fixed shape. However, recent advances have highlighted the potential for deformable particles with engineered mechanical properties.^31,32^ For example, DNA nanotechnology has led to the creation of hinge-like particles with controllable stiffness. Adsorption of deformable particles on membranes is an interesting problem because of the interplay between deformations of the particles and membrane. Additionally, membrane deformations can lead to effective interactions between particles.

In this work, we investigated the equilibrium properties of hinge-like particles adsorbed onto the surface of a planar membrane. We studied the effects of adhesion strength and hinge stiffness on the configurations of isolated particles and on the effective, membrane-mediated interactions between two particles. For isolated filaments, increasing the adhesion strength causes the membrane to deform more strongly around the particles. Figure 2 shows the impact: The membrane has a negligible effect on the configurations of particles when adhesion is weak, but it causes a straightening of the particles when adhesion is stronger. Stiffer hinges require stronger adhesion, and hence larger membrane deformations, to be significantly deformed. Physically, there is a competition between membrane bending energy and hinge deformation energy while maintaining contact contact between the two. With sufficiently strong adhesion, the membrane is significantly deformed, and the particles tend to straighten to decrease the membrane bending energy.

We further investigated interactions between two adsorbed particles. When the adsorption strength is sufficiently large, flexible hinges tend to come together in a tip-to-tip configuration. This suggests an effective, membrane-mediated attraction between them. Using umbrella sampling methods, we calculated the potential of mean force (PMF) as a function of the distance between the centers of mass of two hinges. At weaker adsorption strengths, the effective interaction between two particles is repulsive due to entropic effects. In contrast, at a stronger adsorption strength, two particles experience an effective attraction. For flexible hinges, Figure 4 shows a small energy barrier in the PMF at large distances, followed by an attractive minimum, which is then followed by a strongly repulsive interaction at short distances. The attractive minimum of the PMF is associated with a tip-to-tip configuration. This general shape of the PMF is also observed for the stiff hinge at the same adsorption strength. However, increasing the stiffness of the hinge weakens the attraction, shifts it toward smaller distances, and promotes the occurrence of tip-to-middle configurations. Physically, the attractive interaction is a result of the two particles adopting configurations in which the membrane can more easily deform around them. This results in more pronounced wrapping of the particle that increases the surface area in contact. When the two particles are close together in the strongly repulsive regime, the membrane bending energy decreases, but there is a large, adverse increase in the adhesion energy, which makes these configurations unfavorable.

Deformable particles offer a novel set of design parameters (shape, mechanical properties, regions of compliance, etc.) with which to modulate and potentially control membrane-mediated interactions between them. Our work here demonstrates that varying the strength of adhesion and the stiffness of a deformable hinge changes the effective interactions and can modify the preferred configurations of two interacting particles. Engineering the properties of deformable particles gives a mechanically-tunable means of controlling effective interactions between particles and ultimately their self-assembly on membranes. Interesting future directions include studying many-particle effects, self-assembly, and impacts on the large-scale morphology of membranes and vesicles.

## Acknowledgements

This work was supported by National Science Foundation CAREER Award PHY-1753017.

